# Fungal Bioaerosols at Five Dairy Farms: A Novel Approach to Describe Workers’ Exposure

**DOI:** 10.1101/308825

**Authors:** Hamza Mbareche, Marc Veillette, Guillaume J Bilodeau, Caroline Duchaine

## Abstract

Occupational exposure to harmful bioaerosols in industrial environments is a real treat to the workers. In particular, dairy-farm workers are exposed to high levels of fungal bioaerosols on a daily basis. Associating bioaerosol exposure and health problems is challenging and adequate exposure monitoring is a top priority for aerosol scientists. Using only culture-based tools do not express the overall microbial diversity and underestimate the large spectrum of microbes in bioaerosols and therefore the identification of new airborne etiological agents. The aim of this study was to provide an in-depth characterization of fungal exposure at Eastern Canadian dairy farms using qPCR and next-generation sequencing methods. Concentrations of *Penicillium/Aspergillus* ranged from 4.6 × 10^6^ to 9.4 × 10^6^ gene copies/m^3^ and from 1 × 10^4^ gene copies/m^3^ to 4.8 × 10^5^ gene copies/m^3^ for *Aspergillus fumigatus*. Differences in the diversity profiles of the five dairy farms support the idea that the novel approach identifies a large number of fungal taxa. These variations may be explained by the presence of different and multiple sources of fungal bioaerosols at dairy farms. The presence of a diverse portrait of fungi in air may represent a health risk for workers who are exposed on a daily basis. In some cases, the allergen/infective activity of the fungi may not be known and can increase the risks to workers. The broad spectrum of fungi detected in this study includes many known pathogens and proves that adequate monitoring of bioaerosol exposure is necessary to evaluate and minimize risks.

**Importance:** While bioaerosols are a major concern for public health, accurately assessing human exposure is challenging. Highly contaminated environments, such as agricultural facilities, contain a broad diversity of aerosolized fungi that may impact human health. Effective bioaerosol monitoring is increasingly recognized as a strategic approach for achieving occupational exposure description. Workers exposure to diverse fungal communities is certain, as fungi are ubiquitous in the environments and the presence of potential sources increase their presence in the air. Applying new molecular approaches to describe occupational exposure is a necessary work around the traditional culture approaches and the biases they introduce to such studies. The importance of the newly developed approach can help to prevent worker’s health problems.

## Introduction

Exposure to airborne microbial flora or bioaerosols in the environment, whether from indoor or outdoor sources, is an everyday phenomenon that may lead to a wide range of human diseases. Compared to other well-described microbial habitats, such as water and soil, little is known about the diversity of airborne microbes (1, 2, 3). Whether aerosolized from natural sources (e.g., wind) or human activities (e.g., industrial processes), the dispersal of bioaerosols can impact public health due to the presence of highly diverse and dynamic microbial communities in urban and rural environments. These impacts range from allergies to asthma and can lead to exposure to pathogens (4, 5, 6, 7). Occupational exposure to harmful bioaerosols in industrial environments can be worrisome depending on the types of raw materials present, and the disturbance and the intensity of air movement and ventilation. For example, animal feeding operations involve various sources of biological material potentially associated with respiratory problems (8, 9, 10, 11, 12, 13).

Fungal bioaerosols consist of spores, mycelium fragments and debris which are easily inhaled by workers and cause myriad symptoms including allergies, irritation and opportunistic infections. Long-term lung exposure to fungal bioaerosols can be associated with chronic diseases while the effects of short-term exposure range from irritation of the eyes and nose to coughing and a sore throat (14, 15). Dairy-farm workers are exposed to high levels of fungal bioaerosols on a daily basis. In fact, fungal concentrations in the air at dairy farms were reported to be higher than bacterial concentrations and may reach up to 10^11^ colony-forming units/m^3^ (16). At dairy farms, hay and straw are important sources of fungal bioaerosols, as fungi naturally colonize those substrates, especially if there are high moisture levels (17, 18, 19). Building type and management practices (e.g. free stall, use of various bedding materials, ventilation type) also influence the fungal load in bioaerosols.

The inhalation of large concentrations of fungal bioaerosols can lead to a variety of respiratory problems. The major allergy-related diseases caused by fungi are allergic asthma (20, 21, 22), allergic rhinitis (23, 24), allergic sinusitis (25), bronchopulmonary mycoses (26, 27), and hypersensitivity pneumonitis (28, 29, 30). The latter includes farmer’s lung disease (allergic alveolitis), a disease specific to dairy farm workers (31, 32). Furthermore, a component of the fungal cell wall ((1–3)-β-D glucan), is believed to play a role in pulmonary inflammation, increased sensitivity to endotoxins and pulmonary embolisms (33, 34, 35). Some respiratory symptoms are also associated with fungal exposure including mucous membrane irritation syndrome, nasal congestion, sore throat, and irritation of the nose and eyes (36, 37, 38, 39).

The link between exposure to fungi and occupational diseases is often difficult to prove due to undocumented fungi in bioaerosols. This lack of information is primarily due to the methods used to describe fungi present in the workplace. In diversity studies, culture methods are associated with well-known biases as only the viable/culturable portion of the samples is represented. Using culture-independent molecular methods is a good solution for getting around the non-viable/non-culturable limits of the commonly used culture-based methods. Molecular methods are based on the detection of the genetic material of organisms present in a given sample. Applying these methods to samples from composting and biomethanization environments allowed the identification and quantification of fungal bioaerosols present and a better understanding of human exposure (40, 41). In dairy farms, only culture-dependent methods have been used to assess occupational exposure or ambient fungal aerosols (42, 43, 44, 45). All of the previous studies identified the same frequently encountered genera including *Aspergillus*, *Penicillium, Cladosporium* and *Alternaria*. In Canada, the most recent study that described the airborne fungal microflora in dairy farms is from 1999 (16).

Because of the dearth of information about fungal diversity and concentrations in bioaerosols at dairy farms, the aim of this study was to provide an in-depth characterization of fungal exposure at Eastern Canadian dairy farms using qPCR and next-generation sequencing methods.

## Results

Concentrations of fungal bioaerosols using culture methods to capture the viable spores and qPCR for DNA quantification of *Penicillium*/*Aspergillus* genera and *Aspergillus fumigatus* species are shown in Fig.1. Using culture methods, the results ranged from 3.2 × 10^6^ to 8.2 × 10^6^ CFU/m^3^ in samples from the five dairy farms (DF1 to DF5). A strong correlation was observed between concentrations obtained by culture methods and those obtained by qPCR targeting *Penicillium* and *Aspergillus* (*PenAsp*). Concentrations of *PenAsp* ranged from 4.6 × 10^6^ to 9.4 × 10^6^ gene copies/m^3^ at the five dairy farms. Greater variance was observed in concentrations of *Aspergillus fumigatus* which, ranged from 1 × 10^4^ gene copies/m^3^ at DF3 to 4.8 × 10^5^ gene copies/m^3^ at DF2. Concentrations of *Aspergillus fumigatus* at DF1, DF4 and DF5 were 3 × 10^4^, 2.9 × 10^4^ and 3.9 × 10^5^ gene copies/m^3^, respectively. The highest concentrations of *PenAsp* coincided with the highest concentrations of *Aspergillus fumigatus* as observed at DF2 and DF5 (Fig.1). The gap between the two concentrations was more notable in results from DF1, DF3 and DF4 where concentrations of *Aspergillus fumigatus* were lower.

**Figure 1:**
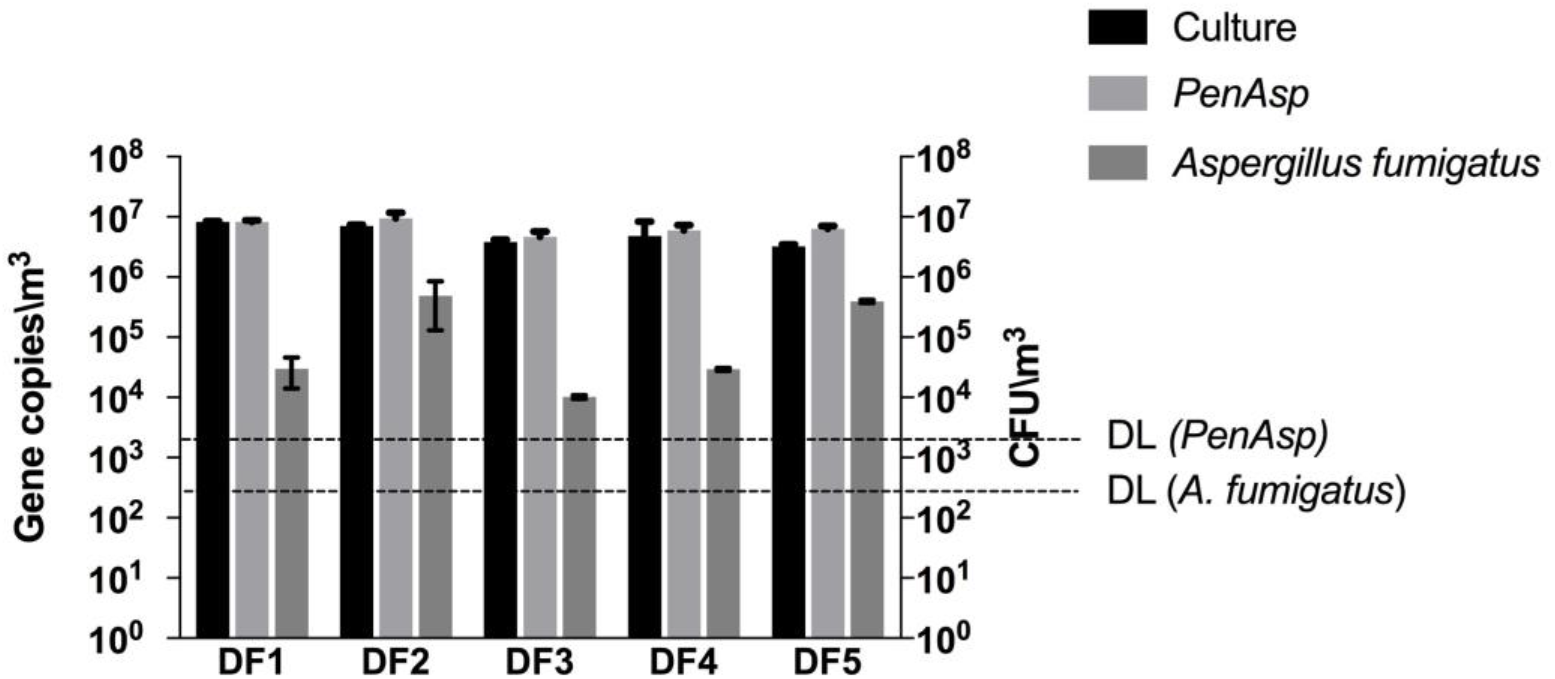
Concentrations of viable spores of mesophilic fungi (from culture), *Penicillium/Aspergillus* (*PenAsp* from qPCR) and *Aspergillus fumigatus* (from qPCR) in air samples collected from five different dairy farms. The detection limit of the qPCR run was 2 × 10^3^ Gene copies/m^3^ for *PenAsp* and 5 × 10^2^ Gene copies/m^3^ for *Aspergillus fumigatus*.

Samples were separate by four categorical variables: *Type of milking*, *animal space, cattle feed*, and *type of ventilation*. Concentrations of *PenAsp* between groups of samples within each of those categories were compared. The same comparison was made for *Aspergillus fumigatus* concentrations. No significant differences (p ≤ 0.05) in concentrations were found between the groups of samples for any of the four variables for either *PenAsp* or *Aspergillus fumigatus* (Table 5).

Fungal communities were described by Illumina Miseq sequencing of the ITS1 region of the fungal ribosomal RNA encoding gene. After quality filtering, dereplication and chimera checking, 307 304 sequences were clustered into 188 OTUs. In order to confirm that the sequencing depth was adequate to describe the fungal diversity at each of the sampling sites, rarefaction analyses were performed using the observed OTUs alpha diversity metric. The lowest-depth sample parameter was used to determine the sequencing depth of the rarefaction analyses which was approximately 40 000 sequences per sample. Samples with a sequencing depth lower than 40 000 were excluded from analyses. The higher the sequencing depth, the more likely it is that the true diversity of the fungi in aerosols is captured. All of the samples from the five dairy farms met this criterion and were included in the analyses. The values shown in Fig.2 were calculated following these steps: ten values from 10 to 40 000 sequences per sample were randomly selected. For each of these values the corresponding number of OTUs observed was noted for all of the samples. The plateaus observed in the five curves shown in Fig.2 indicate an efficient coverage of the fungal diversity, as no more OTUs were observed even with much greater numbers of sequences per sample.

**Figure 2:**
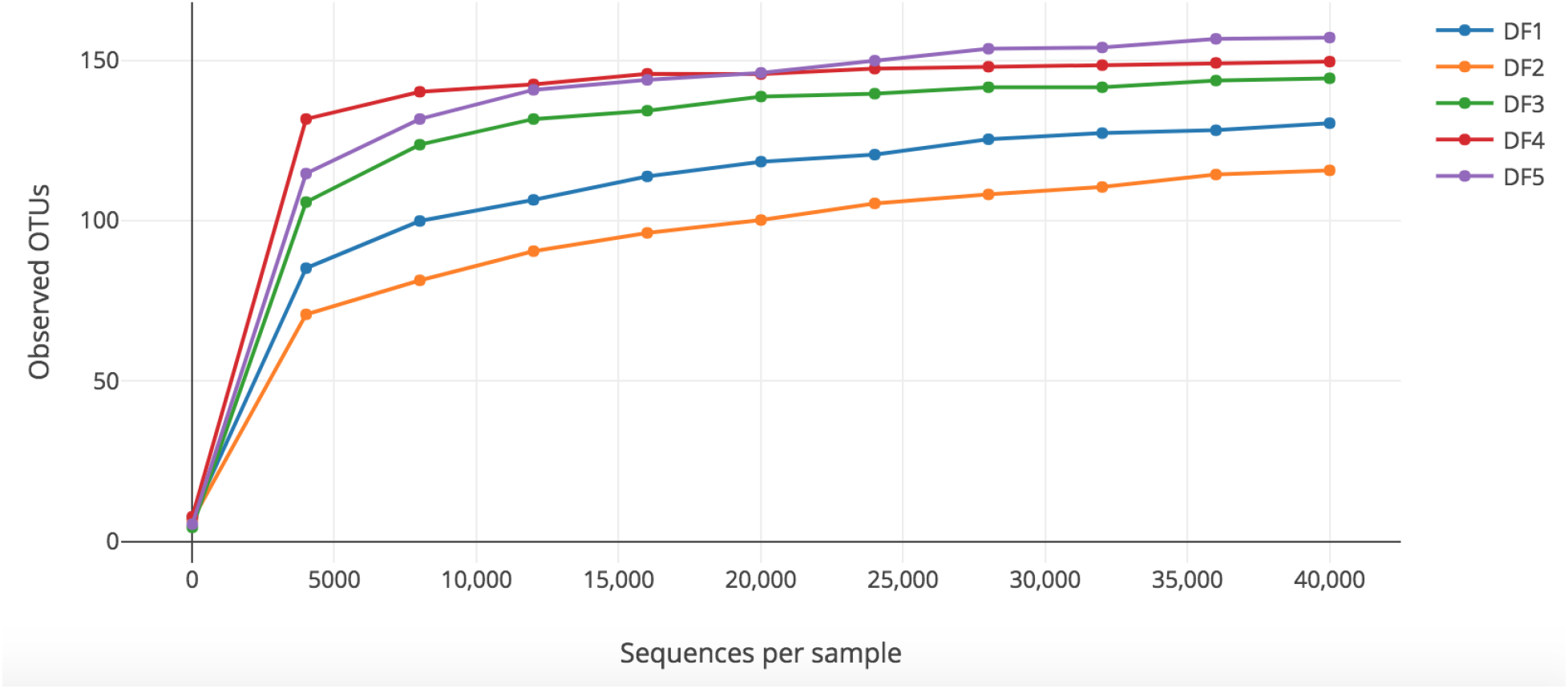
Rarefaction curves obtained from the number of observed OTUs and the sequences per sample for air samples from the five dairy farms visited. The plateau of the curves started at around 5000 sequences.

**Figure 3:**
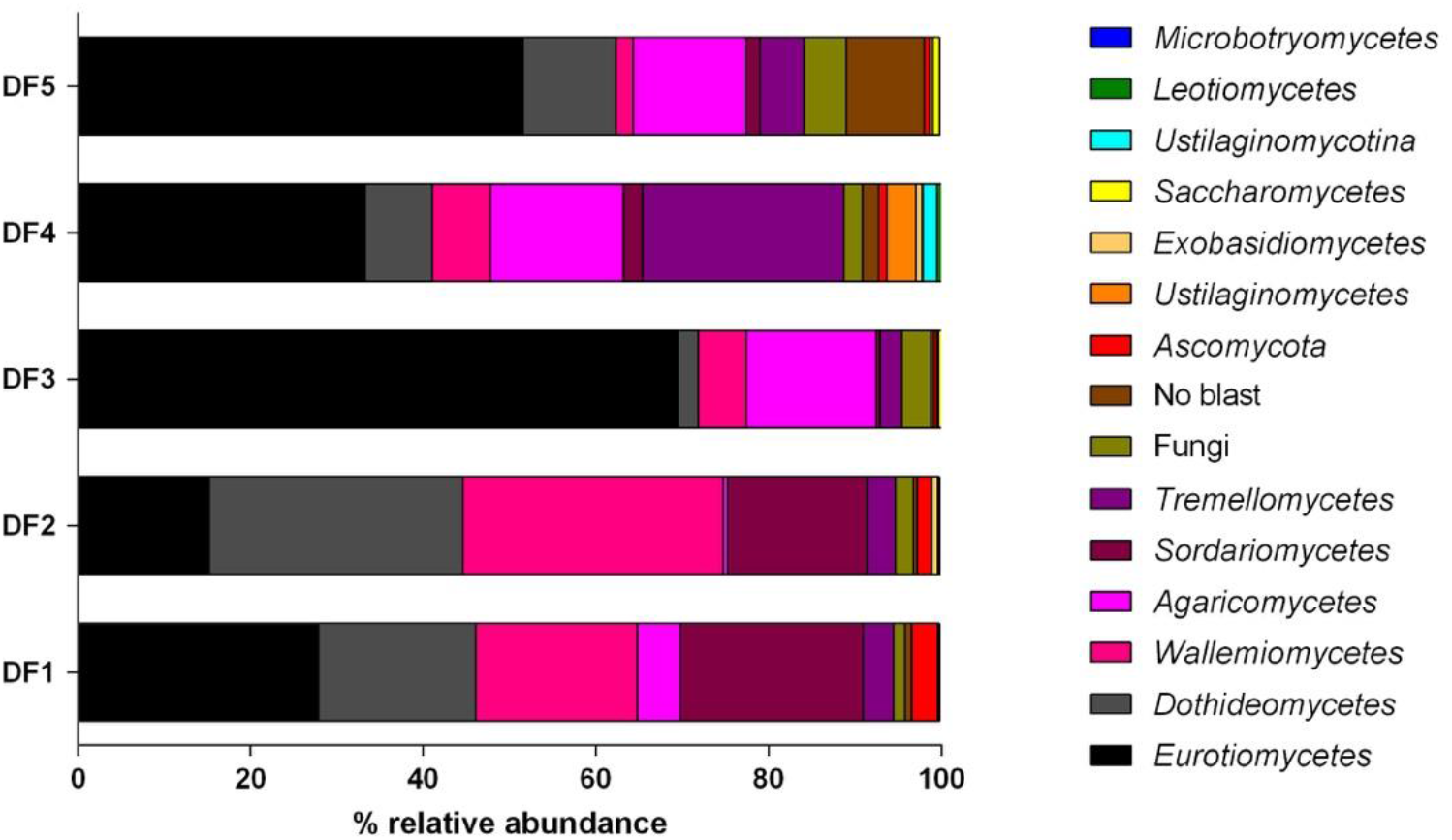
Relative abundance of fungi classes detected in air samples from five dairy farms using next-generation sequencing. While 12 classes were detected, six dominate the diversity profiles. Those six classes have different distributions among the five dairy farms visited.

Many next-generation sequencing surveys of microbial communities aim to compare the composition of different groups of samples (beta diversity). This multivariate approach can be used to assess the effects of several environmental factors on the microbial content of the samples. The environmental factors, or “variables”, are used to separate the samples into different groups. In this case, the same four variables used to categorize qPCR concentrations were also used for the next generation sequencing multivariate analysis and included: *Animal space*, *Cattle feed, Type of milking* and *Type of ventilation*. One of the techniques commonly used by microbial ecologists relies on the creation of a dissimilarity matrix like the Bray-Curtis index. This index was used to evaluate the distance, taken pairwise between samples (representing how closely related samples are). The index uses numbers between 0 and 1, where 0 means the two samples have the same composition and 1 means that they do not share any species. Because the Bray-Curtis dissimilarity matrix uses the absolute abundance of OTUs, it is necessary to use a rarefied OTU table as the input for the dissimilarity calculation. One function of multivariate analyses is to represent inter-sample distances in a 2-dimensional space using ordination (55). To evaluate ordination patterns, one of the most common methods used is the Principal coordinate analyses (PCoA). In this case, the input used for ordination calculation and clustering was the dissimilarity matrix calculated above. The matrix was transformed to coordinates and then plotted using the principal coordinates script in QIIME. Table 6 shows a summary of the results from the PCoA analyses (the PCoA figure is presented as a supplementary file 1). The three principal coordinate axes captured more than 90% of the variation in the DF samples. Samples were coloured according to the four variables to visualize and identify sample clustering. Samples closer to one another are more similar than those that are further away from each other. No obvious sample clustering was observed for any of the four variables. Though they were not clearly clustered, calculations based on *Animal space* and *Cattle feed* were close together than the others. The samples from confined spaces were grouped far from those from the semi-confined space. The forage samples were more closely grouped compared to the samples with concentrates and forage & concentrates combinations. No patterns were observed when samples were coloured according to the *Type of milking* or *Type of ventilation*.

To determine the statistical significance of the variance observed in the PCoA analyses, a PERMANOVA test was performed on the Bray-Curtis dissimilarity matrix. This non-parametric test allows for the analysis of the strength that each variable have in explaining the variations observed between samples (sample clustering). It is based on the ANOVA experimental design but analyzes the variance and determines the significance using permutations, as it is a non-parametric test (56). Whereas ANOVA/MANOVA assumes normal distributions and a Euclidean distance, PERMANOVA can be used with any distance measure as long as it is appropriate to the dataset. The same variables used for color clustering in the PCoA analyses were used with the PERMANOVA test for statistical significance of sample clustering. The QIIME compare categories script was used to generate the statistical results. Results from the PERMANOVA are consistent with the color clustering observations made based on the PCoA analyses. Using a significance of 0.05, the only variables that exhibited significant differences among sample groupings were *Animal space* (p-value = 0.04) and *Cattle feed* (p-value = 0.05). The two other variables tested did not exhibit significant differences (*Type of milking* p-value = 0.61 and *Ventilation* p-value = 0.90).

The taxonomy of the microbes in the air samples collected from the dairy farms was determined by comparing Illumina sequences to the UNITE database. Of the 12 fungal classes detected in samples from the dairy farms (Fig.4) six classes seem to be dominant: *Eurotiomycetes, Dothideomycetes, Wallemiomycetes, Agaricomycetes, Sordariomycetes* and *Tremellomycetes*. However, there is variability in this dominance between the diversity profiles from the five dairy farms. At DF3 and DF5 the class *Eurotiomycetes* have much greater relative abundance than the other classes. In DF2 samples, *Dothideomycetes* and *Wallemiomycetes* are more abundant than the other classes. The *Sordariomycetes* class is particularly more abundant at DF1 compared to the other farms. Fungi from the class *Tremellomycetes* have greater relative abundance at DF4 than any of the other farms. In fact, DF4 has the most diverse profile, in contrast to samples from DF3 where the class *Eurotiomycetes* represents 70% of the relative abundance. *Ustilaginomycotina* were detected only in samples from DF4.

**Figure 4:**
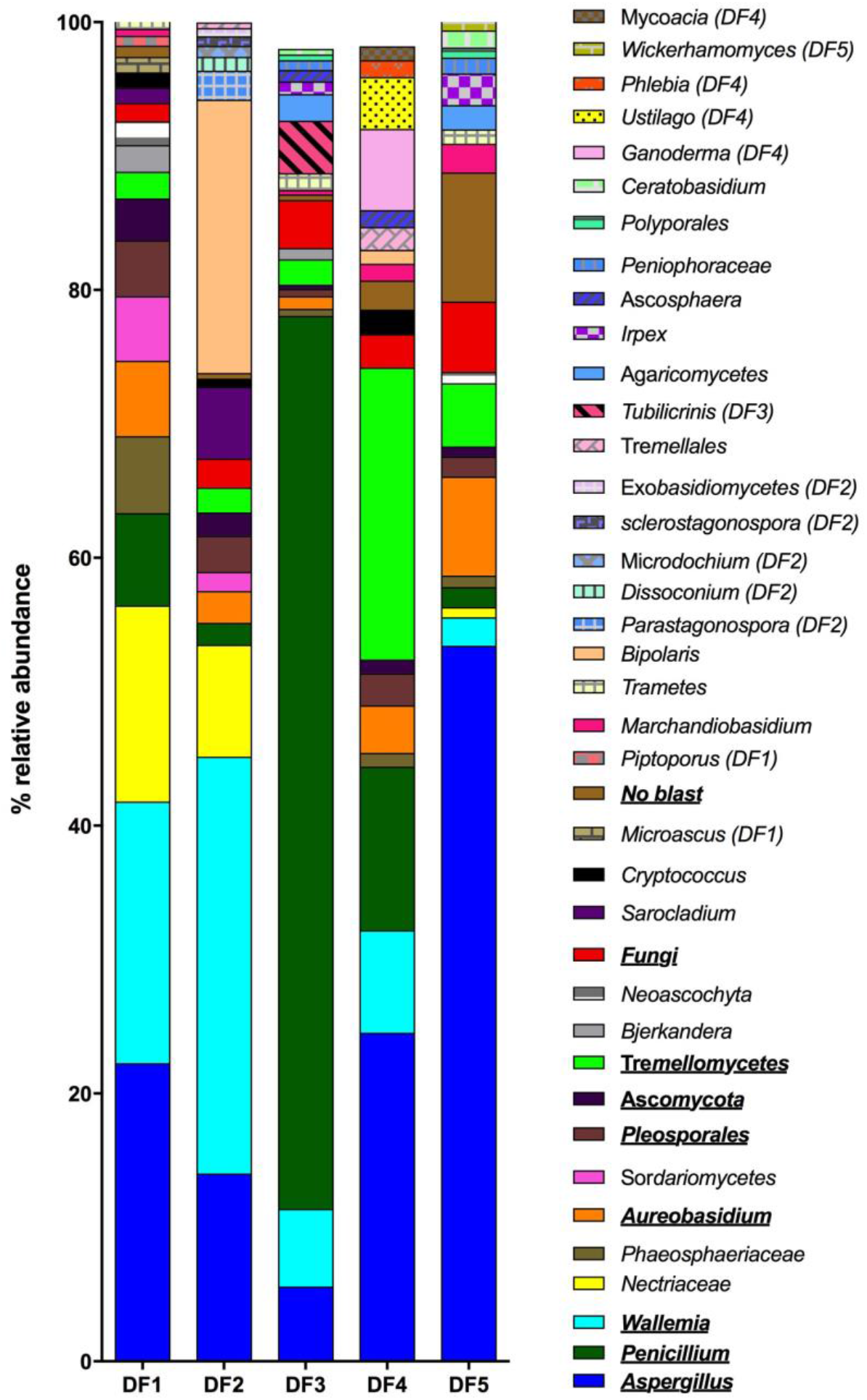
Relative abundance of fungal genera detected in air samples from five dairy farms. The 20 most abundant genera from each dairy farm were included in the analyses. The underlined bold fungi were common in all the dairy farms. The fungi that were detected only in one dairy farm have the DF identification after their name.

Relative abundance of taxa was analyzed more thoroughly by examining the 20 most abundant genera at each dairy farm (Fig.4). From this list, only six fungi were present at all five of the dairy farms: *Aspergillus*, *Penicillium*, *Wallemia, Aureobasidium, Pleosporales* and *Tremellomycetes*. OTUs that were not identifiable to the genus level were identified to the highest taxonomic level (e.g class *Tremellomycetes* and order *Pleosporales*). Similar to observations made based on fungal class, diversity profiles of the genera present were quite variable between the five farms. The least diverse profile was observed in samples from DF3 where *Penicillium* occupied 67% of the abundance. The most diverse profiles were from DF1, DF4 and DF5 as they exhibited the greatest variety of fungal genera. In DF2 samples, 52% of the abundance was made up of *Wallemia* (31 %) and *Bipolaris* (21 %). The diversity profiles from the five dairy farms are larger than what is shown in Fig.4. Due to graphical limitations, only the most abundant fungi are represented. *Piptoporus* and *Microascus* were identified only at DF1. *Exobasidiomycetes*, *Microdochium*, *Dissoconium* and *Parastagonospora* were present at DF2 exclusively*. Tubilicrinis* was detected only at DF3. *Mycoacia, Phlebia, Ustilago* and *Ganoderma* were identified solely at DF4. Finally, *Whickerhamomyces* was specific only to samples from DF5.

The diversity of fungi identified using the culture method was compared with the fungal diversity obtained using next generation sequencing (NGS). Using NGS, fungal genera representing greater than 1% of the total abundance of the five dairy farms combined are presented in Fig.5. For species identified using the culture approach, the fungi identified at more than one dairy farm were grouped together. The fungi that were detected only once by culture were *Trichoderma, Microdochium, Phoma, Apiospora, Botrytis, Conyothirium, Millerozyma, Neosetophoma, Irpex*, and *Debaryomyces*. Those species detected at more than one farm and their relative abundances are presented in Fig.6. The relative abundance of fungi identified by culture was calculated as follows: for each fungus, the number of times that it was isolated from the five dairy farms was calculated. Based on this sum, a percentage of relative abundance was calculated for each fungus and appears in the list in Fig.5. Only four fungi were detected by both approaches: *Penicillium*, *Aspergillus*, *Bipolaris* and *Sarocladium*. Of the 16 fungi isolated using culture techniques, three (*Hyphopichia, Gibellulopsis et Myceliophtora)* were not detected by NGS. The remaining 13, though they do not appear on the list, were detected with a total abundance of less than 1%. Many fungi genera were present but with a total relative abundance of less than 1% making the diversity profile more exhaustive than what is shown in the figures.

**Figure 5:**
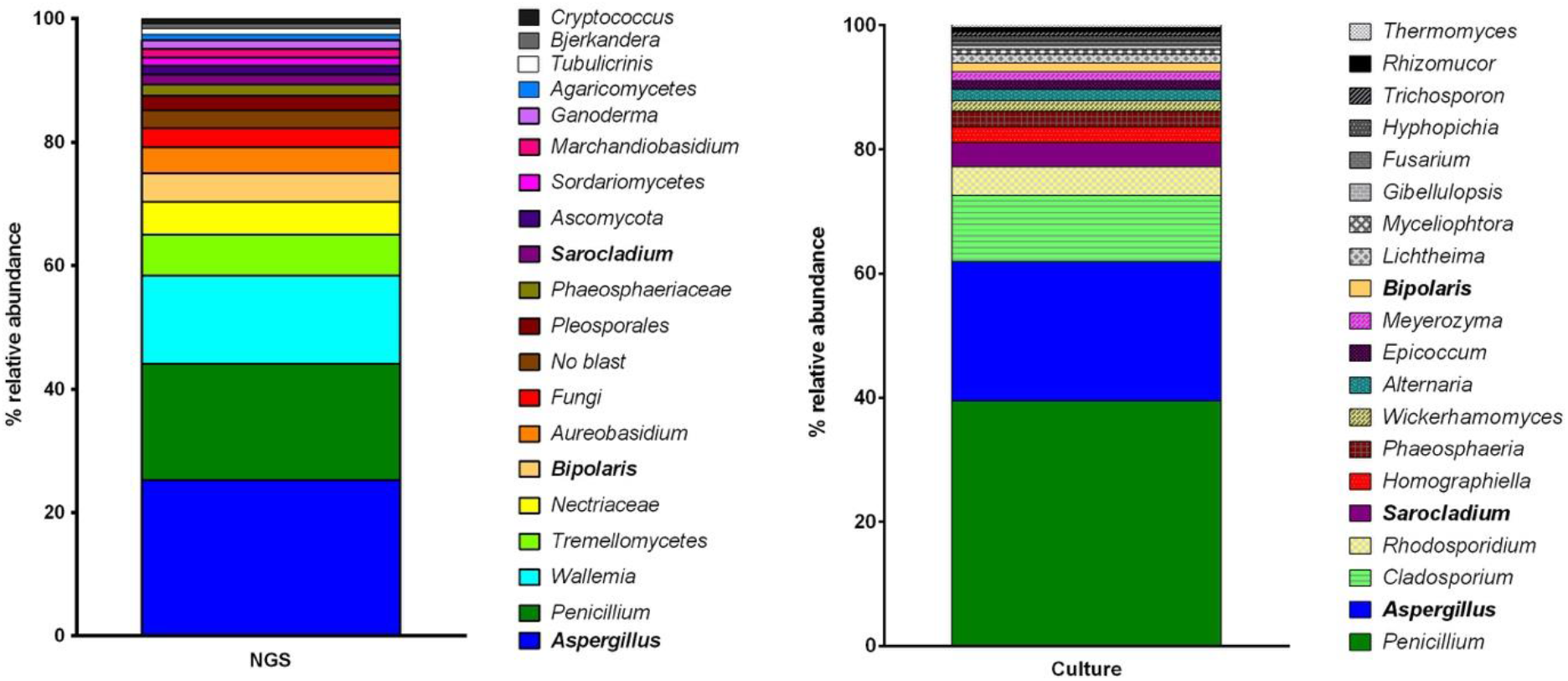
Relative abundance of fungal genera identified by next generation sequencing (NGS) and culture in air samples collected from five dairy farms. Fungi in bold character are common to both approaches.

## Discussion

Using molecular tools has enabled us to provide an in-depth description of the complex biodiversity of bioaerosols in various occupational environments (11, 57, 58, 40, 41). At dairy farms, molecular approaches targeting bacteria allow for a better understanding of the causes of occupational respiratory diseases (59, 60). Since hay and straw are important sources of fungal aerosols and are often ubiquitous at dairy farms, characterizing the fungal diversity of bioaerosols is essential to better understand their role in occupational exposure. As mentioned previously, most studies concerning dairy farm exposure use culture-based methods to study aerosolized fungi. To address the major bias associated with culture methodology, which represents only the viable portion of bioaerosols, this study also used a molecular approach combining qPCR and next generation sequencing to describe the bioaerosol fungal exposure at five dairy farms.

The *PenAsp* qPCR assay is a good indicator of the total quantities of *Aspergillus*, *Penicillium* and *Paecilomyces* conidia in air samples (61). Results of this study showed a strong correlation between concentrations of culturable fungi and *PenAsp*. This correlation supports the idea of using the qPCR *PenAsp* assay as an indicator of total fungal concentration in exposure studies. No significant differences were observed in fungal concentrations obtained from the five dairy farms using qPCR and culture. These concentrations are comparable to concentrations obtained using culture methods almost two decades ago from Eastern Canadian dairy farms (16, 59). This suggests that dairy-farm workers are still at risk for developing diseases linked to fungal exposure. Furthermore, *Aspergillus fumigatus* was specifically quantified in aerosols from areas at dairy farms where humans work because it is a known pathogen that causes aspergillosis, allergic bronchopulmonary aspergillosis and is involved in other pulmonary diseases (62, 63). In some cases, the gap between the concentrations of *PenAsp* and *Aspergillus fumigatus* can be used as an indicator of the diversity of *Aspergillus* and *Penicillium* genera in air samples.

The qPCR analysis allowed the quantification of potentially hazardous fungal spores in bioaerosols. No particular correlation was found between the types of ventilation, animal confinement, cattle feed and milking methods, and concentrations of *PenAsp* and *Aspergillus fumigatus* in aerosols from dairy farms. These results prove that no matter how different the building attributes, animal confinement and types of milking activities are, exposure to fungal bioaerosols should be considered regardless of the modernity of the method used.

The MiSeq Illumina sequencing depth used in this study was adequate for covering the true diversity of fungi in the samples. Targeting the ITS1 genomic region provided an in-depth analysis of the fungal composition of bioaerosols at the five dairy farms. The methodology applied also revealed the variations in fungal communities present in the air (40, 41). Differences in the diversity profiles support the idea that this approach identifies a large number, if not all of the taxa that are responsible for the fungal community changes. These variations in diversity profiles may be explained by the presence of different and multiple sources of fungal bioaerosols at dairy farms. Four variables were chosen to examine these differences more closely. The use of multivariate analyses, PCoA, coupled with a PERMANOVA test, offers a robust statistical significance of sample clustering using distance matrices. Both analyses (PCoA and PERMANOVA) resulted in the same conclusions in regards of sample clustering confirming their usefulness as tools to visualize and measure sample clustering. The main source of the variation in diversity is associated with cattle feed type. Dairy cattle are fed a wide range of feedstuffs, from forage (grasses, legumes, hay, straw, grass silage and corn silage) to concentrates (barley and maize). The presence of *Ustilaginomycotina* and *Exobasidiomycetes* could be explained by the presence of wheat and other grasses. These classes of fungi include the plant pathogen *Tilletia* known to affect various grasses. Biochemical changes in these products, like pH and water content, can affect their fungal composition (64, 65). Animal confinement also affected the fungal composition of bioaerosols. The semi-confined environment consists of an enclosure where dairy cattle have freedom to move around inside the enclosed space. The confined spaces allow no freedom of movement and each cow has its own space. These differences in the density of cows seem to have an impact on the fungal bioaerosols. The type of milking whether automated or manual, and the type of ventilation, either automatic or manual does not seem to have an effect on the fungal content of the bioaerosols collected. However, a limited number of dairy farms were visited during this study and multivariate analyses and sample clustering methods are known to perform better with a large number of samples. A larger number of air samples collected from different dairy farms would be useful to support the findings that milking method and or types of ventilation influences fungal bioaerosol variability. Other factors like building attributes, handling of feed, seed and silage, and method of spreading the bedding can affect the fungal content of the bioaerosols at dairy farms (66, 67, 68, 69). While PCoA gives a cursory assessment of the variables that affect sample clustering, these variables can often be harder to define. A set of chosen explanatory environmental factors does not guarantee that they have true explanatory power. There is always the possibility that an unexplored covariate is the real causal influence on the microbial ecology of the samples (70). Further research including larger sample sizes and additional variables should be conducted.

*Agaricomycetes* are a group of fungi known for their role in wood-decaying activities and in ectomycorrhizal symbiosis (71, 72). The presence of agricultural planting material/products may explain the larger proportions of *Agaricomycetes* identified at DF1 and DF2 compared to the three other farms. Conversely, *Eurotiomycetes* are a class of fungi linked to processes like fermentation used in food processing. Many genera of this class are natural decomposers and are involved in food spoilage (73,74). The presence of natural or processed foods at DF3, DF4 and DF5 might explain the greater abundance of *Eurotiomycetes* detected in the air at those farms. Additionally, the prevalence of *Eurotiomycetes* might also be explained by the presence of silage which is a fermented, high-moisture stored fodder used to feed cattle (75). Members of *Dothideomycetes* and *Tremellomycetes* include several important plant pathogens that grow on wood debris or decaying leaves (76, 77). *Wallemiomycetes* were detected at all five dairy farms. They were most prevalent at DF1 and DF2, representing 20% and 32% of genera detected, respectively. This class includes one order (*Wallemiales*), containing one family (*Wallemiaceae*), which in turn contains one genus (*Wallemia*) (78). These fungi can grow over a wide range of water activity from 0.69 a_w_ to 0.997 a_w_ (79). Water activity is the vapour pressure of water in the product divided by vapour pressure of pure water at the same temperature. High a_w_ support more microbial growth. *Wallemia* have been isolated in air samples from dairy farms in previous studies (80). Airborne *Wallemia* are suspected of playing a role in human allergies like bronchial asthma (81). A study conducted in France identified *Wallemia* as a causative agent of farmer’s lung disease (82). Other prevalent fungal genera commonly found at dairy farms were identified in this study: *Aspergillus*, *Penicillium*, *Cladosporium*, *Alternaria, Nigrospora* and *Periconia*.

For relative abundance, differences observed in the diversity profiles obtained by next generation sequencing (NGS) and culture methods may be explained by the hypothesis that the culture approach may be biased toward fungi from the rare biosphere. These results are consistent with the conclusions made by Shade and his collaborators (83) regarding the complementarity of culture-dependent and culture-independent approaches to studying bacterial diversity. The premise of their study is that culture-dependent methods reveal bacteria from the rare biosphere and provide supplemental information to that obtained using a NGS approach. In the current case, this complementarity is true only for abundance. As mentioned previously, only three fungi were detected exclusively by culture, while more than a hundred fungi were identified by NGS and not by culture. This is consistent with the concept that culture methods may reveal less abundant taxa in an environment while NGS provides a more exhaustive diversity profile. To the best of our knowledge this is the first research to compare both approaches for examining aerosols at dairy farms.

The application of the NGS approach revealed a large fungal diversity profile in bioaerosols released from five dairy farms. The presence of a diverse portrait of fungi in air may represent a health risk for workers who are exposed on a daily basis. In some cases, the allergen/infective activity of the fungi may not be known and can increase the risks to workers. More specifically, the following fungi detected are known allergens and/or are opportunistic pathogens: *Aspergillus, Malassezia, Wallemia, Emericella, Fusarium, Alternaria* and *Candida. Malassezia* causes skin disorders and can lead to invasive infections in immunocompetent individuals (84). *Emericella* is a taxon of teleomorphs related to *Aspergillus*. Species of this group are known agents of chronic granulomatous disease (CGD; 85). *Acremonium* causes fungemia in immunosuppressed patients (86). *Fusarium* species are responsible for a broad range of health problems, from local and systemic infections to allergy-related diseases such as sinusitis, in immunodepressed individuals (87). *Alternaria* is an important allergen related to asthma (88).

## Methodology

### Environmental Field Samples

Air samples were collected from five dairy farms in Eastern Canada during summer 2016. At each farm, a sampling site was designated based on where activities that generate the most bioaerosols took place. The buildings at each farm exhibited differences in building type and characteristics (age, volume, ventilation), number of animals present (cows), methods of milking (automatic or manual) and types of animal feed animal were given. Table 1 presents a description of the sampling sites at each dairy farm. At each sampling site, three air samples were collected during the morning milking activity, when workers are exposed to the most bioaerosols, for a total of 15 samples.

**Table 1:** Description of the sampling sites and the parameters affecting the sampling environments

|  | *Type of milking* | *Animal space* | *Cattle feed* | *Ventilation* | *Temperature* | *Time of sampling* |
| --- | --- | --- | --- | --- | --- | --- |
| **DF1** | manual | confined | forage | natural | 22°C | 6 am – 9 am |
| **DF2** | automatic | confined | forage | mechanical | 21°C | 7 am – 10 am |
| **DF3** | manual | confined | concentrates | natural | 19°C | 7 am – 10 am |
| **DF4** | automatic | confined | concentrates<br>& forage | mechanical | 20°C | 6 am – 9 am |
| **DF5** | manual | semi-confined | forage | mechanical | 23°C | 11 am – 2 pm |

### Air Sampling

A liquid cyclonic impactor Coriolis μ^®^ (Bertin Technologies, Montigny-le-Bretonneux, France) was used for collecting air samples. The sampler was set at 200 L/min for 10 minutes (2m^3^ of air per sample) and placed within 1-2 meters of the source. The air flow in the sampler creates a vortex through which air particles enter the Coriolis cone and are impacted in the liquid. Fifteen millilitres of a phosphate buffer saline (PBS) solution with a concentration of 50 mM and a pH of 7.4 were used to fill the sampling cone.

### Culture-Based Approach to Study Fungal Diversity

One millilitre of the 15ml Coriolis sampling liquid was used to perform a serial dilution from 10^0^ to 10^−4^ concentration/ml. The dilutions were made using 0.9% saline and 0.1% Tween20 solution and were performed in triplicate. Tween20 is a detergent that makes spores less hydrophobic and easier to collect. One hundred microlitres of each triplicate were plated on Rose Bengal Agar with chloramphenicol at a concentration of 50 μg/ml. Half of the petri dishes were incubated at 25°C for mesophilic mould growth and the other half at 50°C for thermophilic mould growth, specifically *Aspergillus fumigatus*. After 5 days of incubation, moulds were identified and counts were translated into CFU/m^3^.

#### Identification of Isolates

Spores from cultured fungi were recovered in one millilitre of a 0.9 % saline and 0.1% Tween20 solution and stored in an Eppendorf tube. Two hundred microlitres of the collection liquid were placed in an FTA card (sample collection card; Qiagen, Mississauga, Ontario, Canada). Five punches from the inoculated zone of the FTA card were placed in a microtube and washed three times with the FTA purification agent. The washing step is mandatory as it allows the removal of the chemical substrates in the FTA card that may alter the subsequent amplification step. Forty-eight microlitres of the master mix solution described in table 2 were placed in each microtube followed by amplification and sequencing of the ITS genomic region. The protocol described by White and his collaborators (46) was performed at the CHU (*Centre hospitalier de l’Université Laval*). The following oligonucleotides were used for the ITS region amplification:

> ITS1: 5’-TCCGTAGGTGAACCTGCGG-3’
>
> ITS4: 5’-TCCTCCGCTTATTGATATGC-3’

**Table 2:** Master mix of the ITS amplification reaction

| Substance | Concentration | Volume |
| --- | --- | --- |
| PCR-mix<br>(IDT, Coralville, États-Unis) | 5x | 10µl |
| MgCl<sub>2</sub><br>(IDT, Coralville, États-Unis) | 25 nM | 6µl |
| Taq DNA polymerase<br>(IDT, Coralville, États-Unis) | 5 µ/µl | 0.5µl |
| Primer ITS1<br>(Promega, Madison, États-Unis) | 100 µM/769 µl IDTE Buffer pH8 | 0.125µl |
| Primer ITS4<br>(Promega, Madison, États-Unis) | 100 µM/1399 µl IDTE Buffer pH8 | 0.125µl |
| dNTP<br>(IDT, Coralville, États-Unis) | 10 nM | 0.5µl |
| ADN | - | 2µl of ADN (5 punch of the FTA-card) |
| H<sub>2</sub>O | - | 30.75µl |

The identification of the isolates was made by comparing the sequences obtained with sequences in the UNITE database.

### Fungal Spore Concentration by Filtration

The following methods are described in detail by Mbareche and his coauthors 2018 (47). Briefly, the 45ml Coriolis suspension was filtered through a 2.5cm polycarbonate membrane (0.2-mm pore size; Millipore) using a vacuum filtration unit. The filters were placed in a 1.5ml Eppendorf tube with 750μl of extraction buffer (bead solution) from a MoBio PowerLyser^®^ Powersoil^®^ Isolation DNA kit (Carlsbad, CA, U.S.A) and a 0.3cm tungsten bead. The filters were flash-frozen by placing the Eppendorf tube in a 99% ethanol solution and dry ice. The frozen filters were then pulverized using the tungsten steel bead in the Eppendorf tube in a bead-beating machine (a Mixer Mill MM301, Retsch, Düsseldorf, Germany) set at a frequency of 20 movements per second for 20 minutes. The liquid containing the pulverized filter particles was used as aliquot for the first step of the DNA extraction procedure.

### DNA Extraction

Using the same bead-beating machine, a second bead-beating step using glass beads at a frequency of 20 movements per second for 10 minutes was performed to ensure that all of the cells were ruptured. Next, a MoBio PowerLyser^®^ Powersoil^®^ Isolation DNA kit (Carlsbad, CA, U.S.A) was used to extract the total genomic DNA from the samples following the manufacturer’s instructions. Next the DNA was eluted in a 100μl buffer and stored at −20°C until subsequent analyses.

### Real-Time PCR Quantification

PCR was performed with a Bio-Rad CFX 96 thermocycler (Bio-Rad Laboratories, Mississauga, CANADA). The PCR mixture contained 2μl of DNA template, 0.150 μM per primer, 0.150 μM probe, and 7.5μl of 2× QuantiTect Probe PCR master mix (QuantiTect Probe PCR kit; Qiagen, Mississauga, Ontario, Canada) in a 15-μl reaction mixture. The results were analyzed using Bio-Rad CFX Manager software version 3.0.1224.1015 (Bio-Rad Laboratories). *Aspergillus fumigatus* was used as positive control and for the standard curves of both qPCR analyses (*Penicillium/Aspergillus* and *Aspergillus fumigatus*). Table 3 presents the primers, probes and PCR protocol used in this study.

**Table 3:**
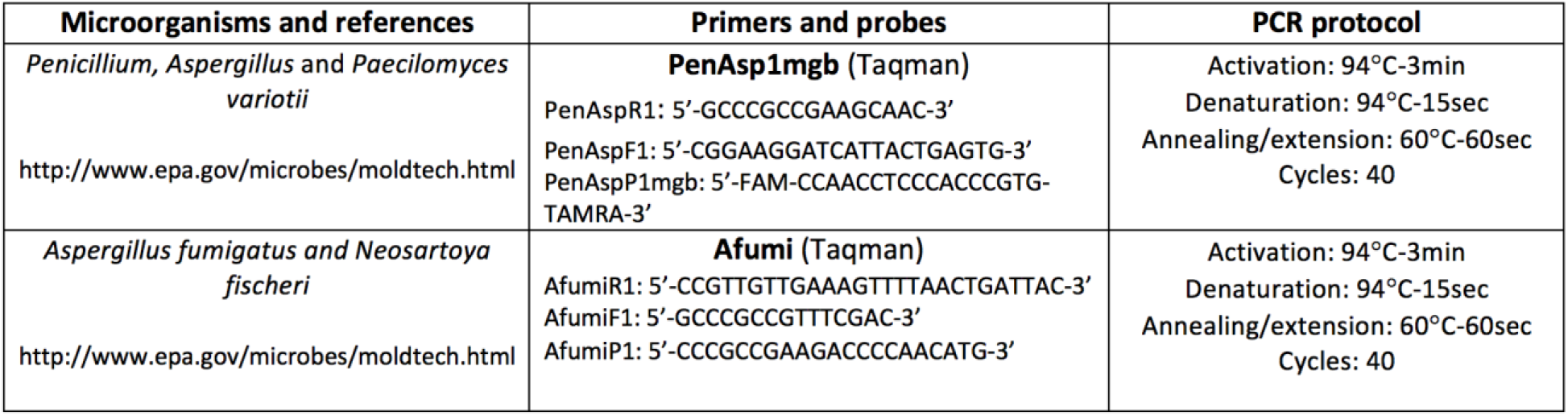
Primers, probes and protocol used for qPCR quantification of selected microorganisms

### Next-Generation Sequencing

The rRNA fungal gene ITS1 was used for the next-generation sequencing analyses. Amplification of the amplicons, equimolar pooling and sequencing were performed at the Plateforme d’analyses génomiques (IBIS, Université Laval, Quebec City, Canada). Briefly, amplification of the ITS1 regions was performed using the sequence specific regions described by Tedersoo et al. (2015) (48) and references therein, using a two-step dual-indexed PCR approach specifically designed for Illumina instruments. First, the gene-specific sequence was fused to the Illumina TruSeq sequencing primers and PCR was carried out on a total volume of 25 μL of liquid made up of 1X Q5 buffer (NEB), 0.25 μM of each primer, 200 μM of each of the dNTPs, 1 U of Q5 High-Fidelity DNA polymerase (NEB) and 1 μL of template cDNA. The PCR started with an initial denaturation at 98°C for 30 s followed by 35 cycles of denaturation at 98°C for 10 s, annealing at 55°C for 10 s, extension at 72°C for 30s and a final extension step at 72°C for 2 min. The PCR reaction was purified using an Axygen PCR cleanup kit (Axygen). Quality of the purified PCR products was verified with electrophoresis (1% agarose gel). Fifty to 100-fold dilution of this purified product was used as a template for a second round of PCR with the goal of adding barcodes (dual-indexed) and missing sequence required for Illumina sequencing. Cycling conditions for the second PCR were identical to the first PCR but with 12 cycles. The PCR reactions were purified as above, checked for quality on a DNA7500 Bioanlayzer chip (Agilent) and then quantified spectrophotometrically with a Nanodrop 1000 (Thermo Fisher Scientific). Barcoded Amplicons were pooled in equimolar concentration (85 ng/μl) for sequencing on the illumina Miseq.

The oligonucleotide sequences used for amplification are presented in Table 4.

Please note that primers used in this work contain Illumina specific sequences protected by intellectual property (Oligonucleotide sequences © 2007-2013 Illumina, Inc. All rights reserved. Derivative works created by Illumina customers are authorized for use with Illumina instruments and products only. All other uses are strictly prohibited.).

**Table 4:**
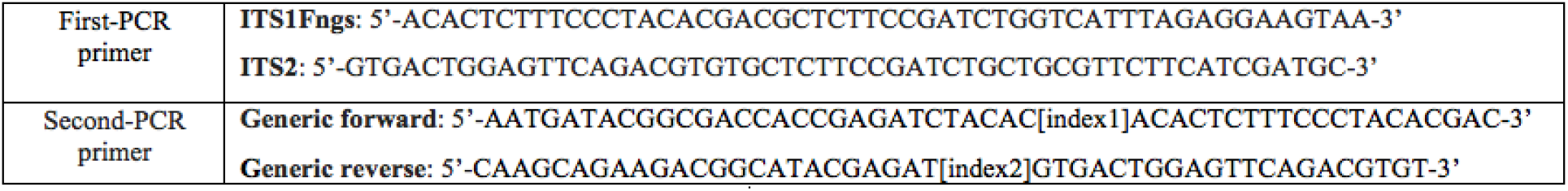
Primers used for Illumina amplification

**Table 5:** Comparison of p-value of the concentrations obtained by qPCR between groups of samples within four environmental factors using Kruskal-Wallis one-way analysis of variance.

| Environmental factors | *PenAsp* (p-value) | *Aspergillus fumigatus* (p-value) |
| --- | --- | --- |
| *Animal Space*<br>(confined vs semi-confined) | 0.09 | 0.06 |
| *Cattle feed*<br>(forage vs concentrates vs concentrates & forage) | 1 | 0.07 |
| *Type of milking*<br>(automatic vs manual) | 0.74 | 0.07 |
| *Type of Ventilation*<br>(mechanical vs natural) | 0.8 | 0.08 |

**Table 6:** Summary of the parameters and results of the principal coordinates analysis of air samples collected from five dairy farms including the statistical significance of the sample clustering. The PCoA was calculated using the Bray-Curtis dissimilarity based on ITS1 sequences. The three principal coordinate axes captured over 90% of the variation in the input DF samples. The statistical significance of the variation observed with the PCoA analyses was determined using a PERMANOVA statistical test. Four environmental variables were used for sample clustering, only two were statistically significant (*Animal space* and *Cattle feed*).

| Environmental factors | Sample clustering | Permanova (p-value) |
| --- | --- | --- |
| *Animal Space*<br>(confined vs semi-confined) | √ | 0.04 |
| *Cattle feed*<br>(forage vs concentrates vs concentrates & forage) | √ | 0.05 |
| *Type of milking*<br>(automatic vs manual) | X | 0.61 |
| *Type of Ventilation*<br>(mechanical vs natural) | X | 0.90 |

### Bioinformatics Workflow

The bioinformatics workflow used in this study was developed during a compost study by Mbareche et al. (40). Briefly, after demultiplexing the raw FASTQ files, the reads generated from the paired end sequencing using Mothur v 1.35.1 were combined (49). Quality filtering was performed using Mothur by discarding reads with ambiguous sequences. Reads shorter than 100 bp and longer than 450 bp were also discarded. Similar sequences were combined to reduce the computational burden, and the number of copies of the same sequence was displayed. This dereplication step was performed using USEARCH version 7.0.1090 (50). The selected region of fungal origin was then extracted from the sequences with ITSx which uses HMMER3 (51) to compare input sequences against a set of models built from a number of different ITS region sequences found in various organisms. Only the sequences belonging to *fungi* were kept for further analyses. Operational taxonomic units (OTUs) with a 97% similarity cut-off were clustered using UPARSE 7.1 (52). UCHIME was used to identify and remove chimeric sequences (53). QIIME version 1.9.1 (54) was used to assign taxonomy to OTUs based on the UNITE fungal ITS reference training data set for taxonomic assignment and to generate an OTU table. The fungal diversity analysis was achieved by using different QIIME scripts. The alpha and beta diversity scripts used are listed in the following link: http://qiime.org/scripts/.

### Statistical Analyses

Concentrations of *PenAsp* were compared with the Kruskal-Wallis one-way analysis of variance. The test was performed using the software R version 3.3.2 with RStudio Version 0.99.486. The same analysis was performed comparing concentrations of *Aspergillus fumigatus*.

To determine the statistical significance of the variation observed with the multivariate analyses (PCoA), a PERMANOVA test was performed on the Bray-Curtis dissimilarity matrix. The *compare categories* QIIME script was used to generate the statistical results. Because PERMANOVA is a non-parametric test, significance is determined through permutations. The number of permutations used is 999. P-value ≤ 0.05 was considered statistically significant. Detailed information about the performance of the test are presented in the multivariate section of the results.

## Conclusion

Bioaerosols from Eastern Canadian dairy farms contain high concentrations of highly diverse fungi. This study demonstrated that fungal bioaerosols have large diversity profiles. It also adds another piece to the puzzle regarding the complexity of bioaerosols relative to the sources present. This study also highlights the importance of using a high-throughput sequencing method combined with real-time PCR assay for quantification and an in-depth characterization of fungal diversity in bioaerosols to better assess occupational exposure. As air samples were collected during activities where workers are present, this study shows that human exposure to harmful fungi may be higher during milking activities (automatic or manual), as well as during handling of feed and silage and when spreading bedding. Based on the results of this investigation, the authors strongly recommend taking action to reduce workers’ exposure to bioaerosols. Such measures include increased air-exchange rates, better confinement and source ventilation. If these measures cannot be applied, we recommend skin and respiratory protection for workers who are exposed on a daily basis as a means to reduce continuous exposure to harmful fungi present in bioaerosols. The broad spectrum of fungi detected in this study includes many know pathogens and proves that adequate monitoring of bioaerosol exposure is necessary to evaluate and minimize risks.

## ACKNOWLEDGEMENTS

HM is a recipient of the FRQNT PhD scholarship and received a short internship scholarship from the Quebec Respiratory Health Network. This work was supported by the Institut de Recherche Robert-Sauvé en Santé et en Sécurité du Travail (2014-0057). We are grateful to all of the employees of the dairy farms that participated in this study. We are also grateful to Julien Duchaine and Philippe Bercier that were in the field for their technical assistance. The authors are thankful to Amanda Kate Toperoff and Michi Waygood for English revision of the manuscript. CD is the head of the Bioaerosols and respiratory viruses strategic group of the Quebec Respiratory Health Network.

